# Behavioral ontogeny in a pelagic tunicate reveals the deep origins of chordate behavioral developmental plasticity

**DOI:** 10.64898/2025.12.03.692109

**Authors:** Oleg Tolstenkov, Rodolfo da Silva Mazzarini Baldinotti, Sissel Norland, Anne Elin Aasjord, Daniel Chourrout, Marios Chatzigeorgiou

**Affiliations:** Michael Sars Centre, Faculty of Science and Technology, University of Bergen, Norway

## Abstract

Behavioral developmental plasticity refers to the lasting changes in an organism’s behavior that occur during development in response to external and internal cues, enhancing survival and reproduction. While well studied in vertebrates, its occurrence in other chordates, including tunicates, the closest relatives of vertebrates, remains unclear.

We built a behavioral atlas of the planktonic tunicate *Oikopleura dioica* across its life cyclefrom larval to adult stages using high-throughput tracking and unsupervised analysis. Swimming kinematics, exploration, and thigmotaxis increased with development. Adults exhibit a unique innovation, known as the house, a complex extracellular filtration structure encasing the animal trunk, which drastically alters locomotion and exploration compared to free-swimming individuals. Postural variation during ontogeny is captured by seven basic shapes (“eigenoikopleuras”), with complex shapes becoming more frequent over time. Spatiotemporal embedding and Hidden Markov modeling revealed that behavioral refinement arises from changes in motor modules and their transitions. House occupancy similarly shifts these modules, indicating context-specific specialization.

This continuous behavioral maturation parallels *Oikopleura*’s neotenic morphology, retaining a larval body plan unlike other tunicates. Our atlas provides a quantitative framework for exploring the evolution of behavioral developmental plasticity in chordates.

## INTRODUCTION

Behavioral development is a complex process, influenced by an animal’s genetic traits, early experiences, ongoing interactions and various environmental factors^1,2^. We are most familiar with developmental plasticity of the behavioral repertoire of our own species (*Homo sapiens*), where certain pivotal changes with far-reaching impact, termed behavioral cusps (e.g. the ability to crawl, walk, speak) dramatically alter our interaction with our environment and other humans ^3,4^.

Outside of humans, the process of behavioral developmental plasticity across large parts (or the entirety) of ontogeny has been studied in a limited number of organisms including the roundworm *Caenorhabditis elegans*^5^, the freshwater fish *Danio rerio*^6,7^ and *Poecilia formosa* (Amazon mollies)^8^. These animals show surprising behavioral plasticity despite the conservative changes that take place in the nervous system during their post-embryonic ontogeny^9–12^.

These studies cover a minute fraction of the animal diversity on our planet, therefore there is a significant gap in our understanding of behavioral developmental plasticity across most branches of the phylogenetic tree of life. Vertebrates belong to the phylum Chordata which is composed of three groups (cephalochordates, tunicates and vertebrates). The discovery that cephalochordates are basal within chordates and thus the tunicates are the sister group of vertebrates, suggested that the adult ancestral chordate animals were free-swimming organisms in contrast to the ascidians which have a freely swimming larva^13^ and a sessile filter-feeding (yet behaving) adult^14^. This finding brought renewed interest in appendicularians, the only tunicates that retain a freely swimming, chordate body plan throughout their life^15,16^ (Figure 1A), likely representing the ancestral tunicate condition, considering their position as the sister group to all other tunicates ^17–20^.

**Figure 1.**
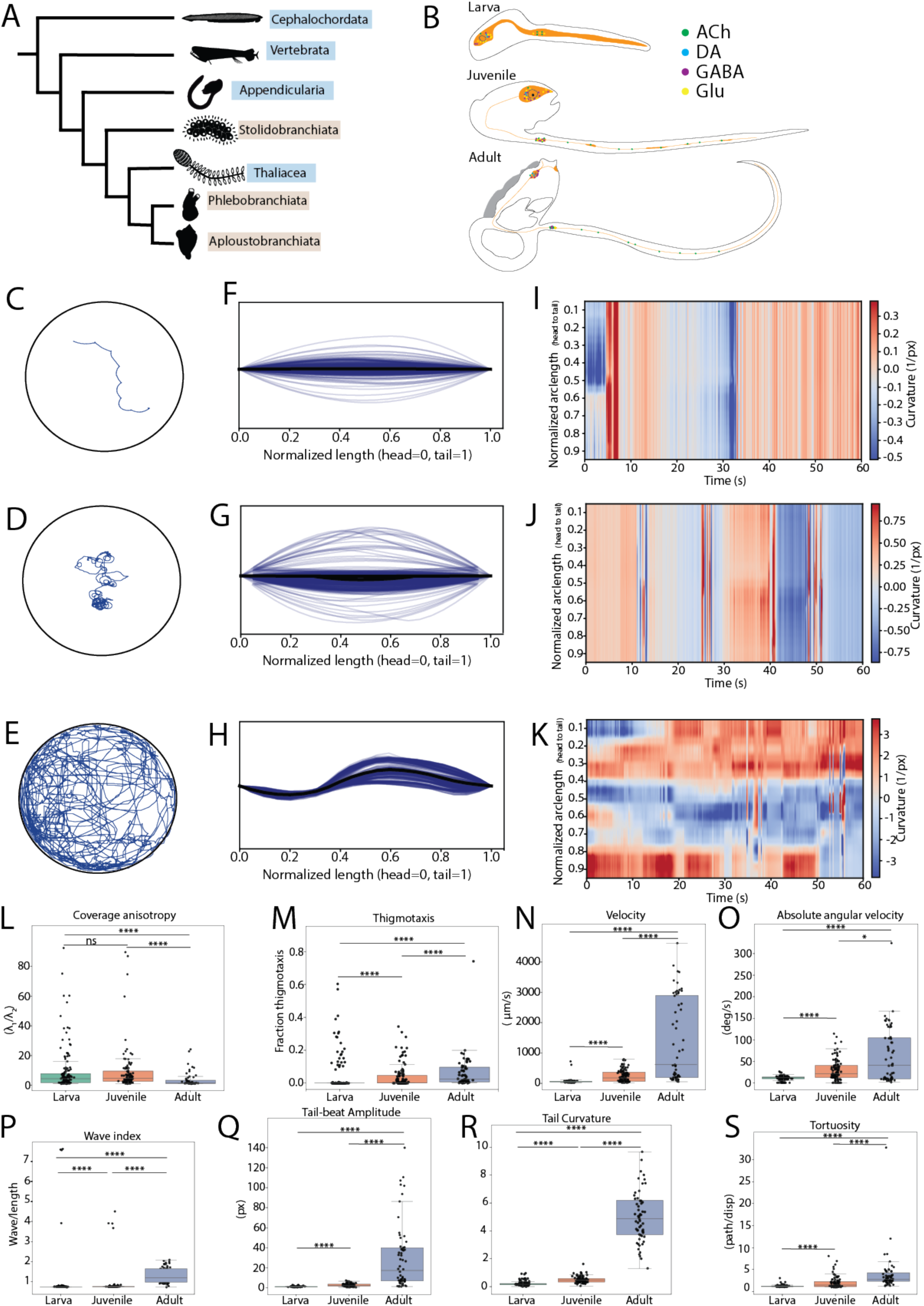
Developmental changes in *Oikopleura doica* locomotor behavior. (A) Phylogenetic position of Appendicularia within chordates. Note that basal groups are the free-swimming cephalochordates within chordates and the free-swimming appendicularians within tunicates. (B) Developmental stages analyzed: larva (top), juvenile (middle), and adult (bottom) with neurotransmitter distributions indicated (ACh, acetylcholine; DA, dopamine; GABA, γ-aminobutyric acid; Glu, glutamate). Scale bars represent body proportions. (C–E) Representative 60 s trajectory traces. (C) Larval stage showing minimal spatial exploration. (D) Juvenile stage displaying increased spatial coverage. (E) Adult stage exhibiting extensive exploratory behavior with complex path structure. (F–H) Normalized centerline overlays (n=200 traces per stage; gray=individual centerlines, dark blue=mean). (F) Larva: extended, relatively straight body. (G) Juvenile: moderate body curvature. (H) Adult: pronounced mid-body bend. Head positioned at (0,0), tail at (1,0) after normalization. (I–K) Spatiotemporal curvature heatmaps over 60 s. Y-axis: normalized body length (0=head, 1=tail); x-axis: time; color: local curvature (1/px). (I) Larva: minimal curvature variation. (J) Juvenile: increased mid-body curvature dynamics. (K) Adult: sustained high curvature in posterior region, reflecting increased amplitude and curvature of tail-beating. (L–S) Developmental comparison of behavioral metrics (individual animals shown as points). Boxplots: median (center line), interquartile range (box), 1.5× IQR (whiskers). Kruskal-Wallis tests with Holm-corrected post-hoc Mann-Whitney U tests: *p<0.05, **p<0.01, ***p<0.001, ****p<0.0001. (L) Coverage anisotropy (dimensionless): larvae exhibit minimal directionality; juveniles and adults show similar moderate anisotropy. (M) Thigmotaxis (fraction of time near boundaries): larvae show minimal boundary preference; adults display strongest thigmotaxis. (N) Velocity (µm/s): adults display 10-fold higher velocities than larvae/juveniles (larvae median=86 µm/s, juvenile=342 µm/s, adult=2,847 µm/s). (O) Absolute angular velocity (deg/s): progressive increase across development (larva=21 deg/s, juvenile=38 deg/s, adult=102 deg/s). (P) Wave index (dimensionless metric of undulatory motion): adults show reduced wave propagation compared to juveniles, consistent with localized tail-beating. (Q) Tail-beat amplitude (px): adults exhibit 5-fold larger amplitudes than earlier stages. (R) Tail curvature (1/px): adults display dramatically increased curvature compared to larvae/juveniles. (S) Tortuosity (path length/displacement, dimensionless): adults show highest path complexity (median=3.2) compared to larvae (1.8) and juveniles (2.1).

The appendicularian *Oikopleura doica* develops in an extremely rapid manner, attaining a motile larval stage within ∼7 hours post fertilization (Figure 1B). This stage is succeeded by a juvenile stage, just prior to which the tail undergoes a remarkable 90 counterclockwise rotation relative to the trunk^15,21,22^ (Figure 1B). This rotation reflects in the direction of the tail beat. In contrast to ascidians, cephalochordates and lower vertebrates whose tails beat in a left-right direction, *Oikopleura* juvenile tail beats in a dorsal-ventral direction^15^. Subsequently, *Oikopleura* matures to the adult form lasting up to 7 days depending on the cultivation temperature (Figure 1B). During adulthood, it lives within a complex mucus structure known as the house ^23^ which it uses for filter feeding through a sequence of precisely controlled behavioral actions^24–26^. The house is replaced multiple times each day when the discarded house is usually clogged ^27,28^.

The development of the *Oikopleura* nervous system across ontogeny is well characterized. Most neuronal cell divisions are complete by the onset of the juvenile stage and cell numbers change only modestly thereafter (60 to 78 neurons in the brain ganglion and stable in the nerve cord, specifically 25-30 neurons in the caudal ganglion and 30 in the caudal nerve cord)^9–11^. Muscle innervation begins at the larval stage, while more complex innervation of muscles has been documented at the adult stage^10^.

Tail movements arise as brief myogenic twitches in post-hatching larvae and consolidate into coordinated tailbeats by the end of the larval stage^26^. Recent studies have begun to describe Oikopleura behaviors related to house building (e.g. nodding) and house function in feeding^24–26,29,30^.

Therefore, *Oikopleura* presents itself as an appealing organism to study behavioral developmental plasticity throughout ontogeny in a tunicate whose swimming is not abolished by the process of metamorphosis^31^.

Here we built a behavioral atlas of *Oikopleura dioica* across its life cycle using high throughput tracking and unsupervised (machine learning based) analysis. We found that kinematic parameters increased during development, as did arena exploration. Kinematics and arena exploration differed markedly depending on whether *Oikopleura* was inside the house (filter-feeding) or outside the house (the period that the animal is freely swimming before inflating a new house). By employing dimensionality reduction, we derived lower dimensional representations of body postures, which we termed “eigenoikopleuras”. With these, we can desribe the majority of postural variance in the *Oikopleura* larvae, juveniles and adults. Leveraging a stage-balanced t-SNE (t-Distributed Stochastic Neighbor Embedding) space, we quantified behavioral dynamics across development, identifying a core set of clusters conserved across development as well as clusters that are stage specific. Similarly, a small subset of clusters distinguishes between in house and out of house behaviors. Finally, we modelled *Oikopleura* behavior using a Hidden Markov model (HMM) revealing the gradual temporal refinement in behavioural output rather than abrupt reorganization.

## RESULTS

To study behavioral development in the appendicularian *Oikopleura dioica* we recorded videos of larvae, juveniles and adults (inside or outside the house) (Movies S1-S4) in our tracking setups^13,32^ and used DeepLabCut^33^ to extract, in a high-throughput manner positional coordinates of multiple body parts of each *Oikopleura* larva, juvenile and adult. Center of mass trajectories suggested that larvae exhibited simple episodic swimming with few changes in direction of travel (Figure 1C). Juveniles exhibited more complex swimming patterns with local runs and turns, yet they explored only a fraction of the arena (Figure 1D). Finally, adults exhibited very complex trajectories exploring most of the arena (Figure 1E).

Larvae and juveniles were characterized by straight midlines which showed small deflections to the right and the left potentially indicating that simple swinging of the tail was sufficient for the animals to move (Figure 1F and 1G; Figure 1I and 1J). In contrast adults were marked by sigmoidal midline skeletons, showing large increase tail curvature, increased tail amplitude and curvature and more waves traveling along their tail compared to larvae and juveniles (Figure 1H-1K).

In line with the changes in swimming and exploratory characteristics, this visual difference in the extent to which they explored the arena was quantified through the arena coverage anisotropy with the larvae showing the highest anisotropy (i.e. only exploring small regions of the arena) and adults exhibiting much lower anisotropy values indicative of exploring the arena in a more thorough and uniform manner (Figure 1L).

Adults displayed a significantly higher thigmotaxis compared to larvae and juveniles (Figure 1M). We have previously shown that *Ciona intestinalis* larvae are also thigmotactic^32^ suggesting that thigmotaxis may be a conserved behavior amongst tunicates.

In addition, behavioral traits such as velocity, angular velocity, tail amplitude, tail curvature and wave index increased progressively across developmental stages, with the most significant increase between the juvenile to the adult stages (Figure 1N-1R). We also quantified the tortuosity (complexity of a trajectory) of the animal trajectories. We found that tortuosity significantly increased during juvenile and adult stages (Figure 1S).

*Oikopleura* maintains efficient filter-feeding capacity by a house that surrounds the animal (Figure 2A), which is secreted from the trunk epithelium in the free-swimming animal (Figure 2B). We asked how the presence of the house modified the behavior of adult *Oikopleura*. We found that while in the house *Oikopleura* adults displayed limited exploration of the arena in contrast to the animals swimming outside the house (Figure 2C and 2D). In line with this, pronounced differences were observed in tail swinging patterns and curvature between inside house to that of a outside house phenotypes (Figure 2E-2H). Animals found outside the house had significantly reduced anisotropy and were more thigmotactic compared to animals inside the house (Figure 2I-2J). We then compared the kinematics of tail movement inside versus outside the house. Compared to inside house animals, we observed animals outside the house had significantly lower tail beating symmetry, wave index and tail curvature in addition to increased tail beating amplitudes (Figure 2K-N). Despite showing limited overall displacement while inside the *house Oikopleura* moves its tail creating a current that draws water through the house’s filters to allow the animal to feed^24,27^. Tail beating inside the house shows greater lateral bias, suggesting that *Oikopleura* adults in the house consistently move their tail more frequently to one side. Tail movements inside the house are characterized by a higher wave index (i.e. the tail has higher curvature but lower tail-beat amplitude). The difference may reflect a behavioral adaptation for feeding in the house. In contrast the symmetric, higher amplitude tail beating of freely swimming adults is more suited for straight trajectory swimming outside the house.

**Figure 2.**
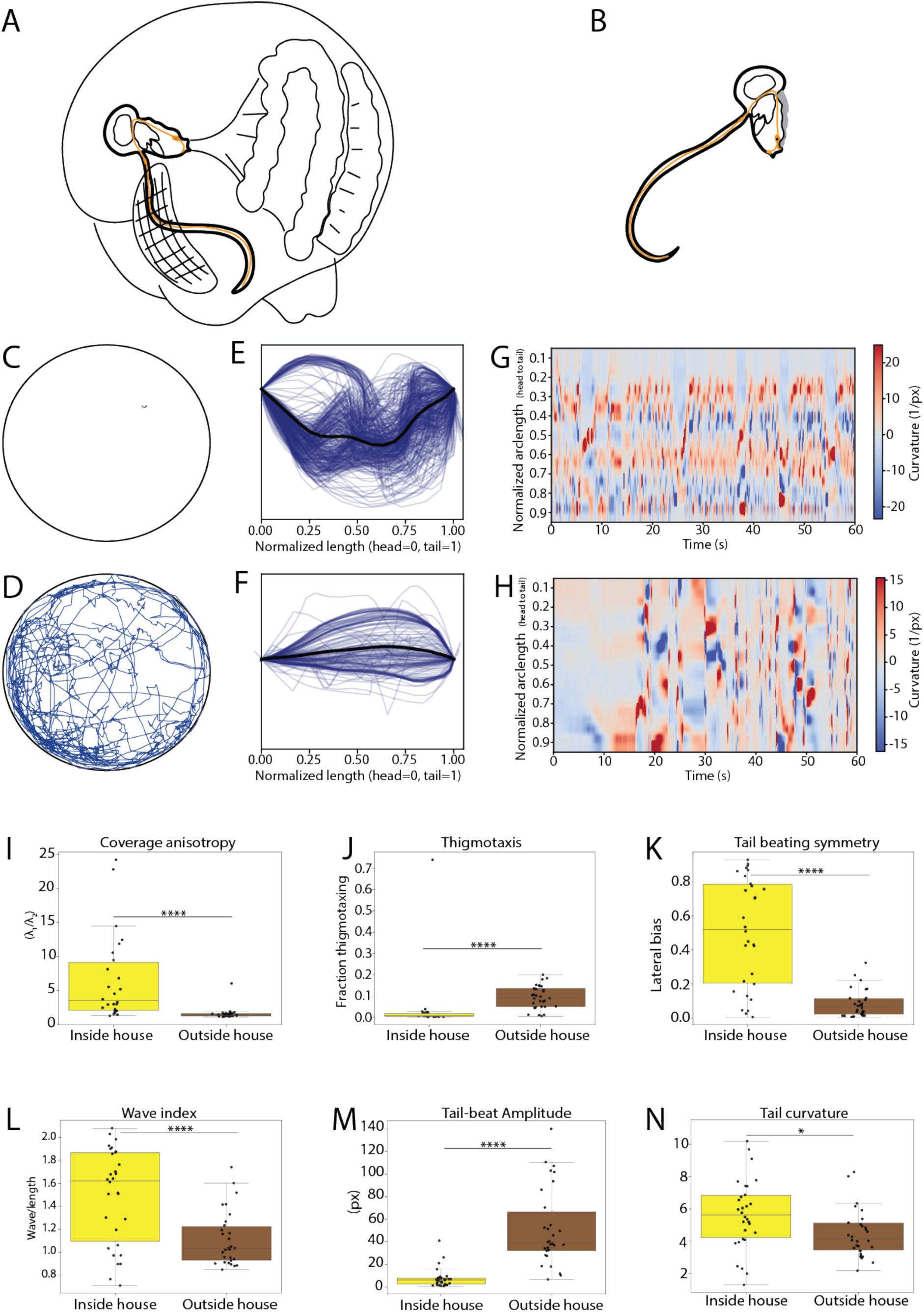
Adult behavior inside versus outside the house. (A) Schematic of adult *Oikopleura* within house structure and free-swimming outside house (B) (please see also Videos S3 and S4). (C–D) Representative 60 s trajectory traces. (C) Outside house: minimal spatial displacement. (D) Inside house: extensive spatial exploration within house boundaries. (E) Normalized centerline overlay inside the house (black=individual, dark blue=mean). (F) Normalized centerline overlay outside house. (G–H) Spatiotemporal curvature dynamics over 60 s. (G) Inside and (H) outside the house. (H) (I–N) Behavioral comparison between contexts (individual animals shown as points). Statistics: Mann-Whitney U tests with Holm correction. Boxplots as in Figure 1. (I) Coverage anisotropy; (J) Thigmotaxis; (K) Tail-beating symmetry (lateral bias, dimensionless); Inside-house animals display strongly biased tail-beating. (L) Wave index. (M) Tail-beat amplitude. Outside-house animals show 6-fold higher amplitudes. (N) Tail curvature.

Overall, our data suggest that *Oikopleura* adults show significant plasticity in their locomotion kinematics and arena exploration depending on whether they are in the house or they have escaped the house and are freely swimming. These results reveal context-dependent behavioral specialization: inside-house animals exhibit restrained, asymmetric movements optimized for filter-feeding, while outside-house animals engage in large-amplitude, symmetric propulsion for navigation.

Features like tail curvature, tail beating symmetry, velocity and others can describe *Oikopleura* postures accurately, however, the richness in these descriptions comes at the cost of a very high dimensionality. We sought to obtain a simpler representation that would describe the broad range of postures that *Oikopleura* can obtain (as represented by a randomly sampled set of curvature values) without losing significant information^13,34,35^. We performed a principal components analysis (PCA) and found that 95% of the observed variance in the curvature data can be explained with 7 eigenvectors (here after called eigenoikopleuras) (Figure 3A). For any given frame in the videos we recorded across all three stages can be approximated as a linear combination of the 7 eigenoikopleuras (PC1-PC7). PC1 captures overall body bending, PC2 reflects an anterior-posterior curvature gradient, and PCs 3-7 encode higher-order shape modes and local curvature features (Figure 3B and 3C).

**Figure 3.**
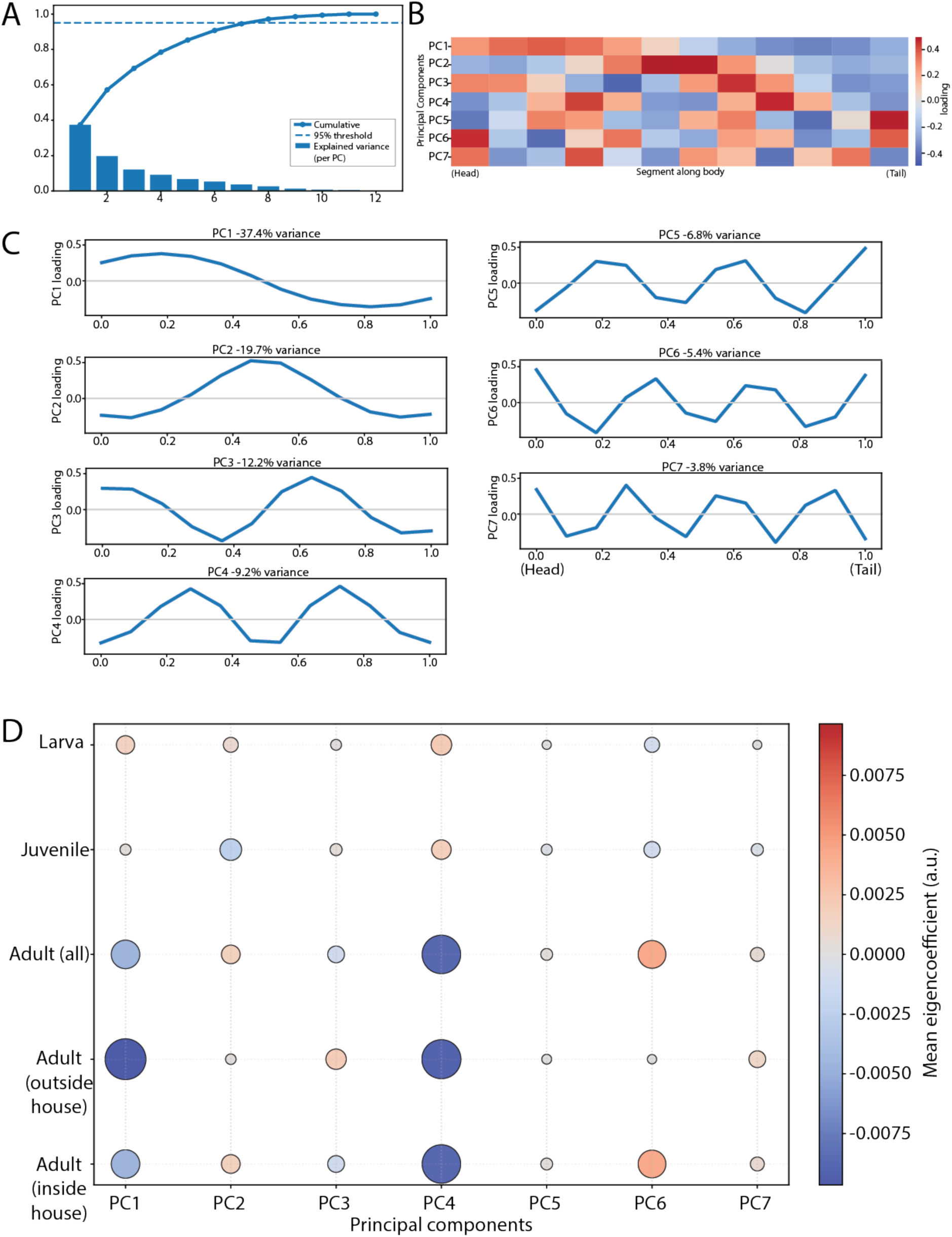
Principal component analysis reveals behavioral mode structure. (A) PCA variance decomposition. Bar plot: variance explained per PC. Line: cumulative variance (7 PCs explain 95.3% total variance). Dashed line: 95% threshold. Seven PCs were retained for subsequent analyses (see Methods). (B) PC loadings heatmap along body axis. Rows: PC1–PC7; columns: 21 evenly spaced points from head (left) to tail (right). Color: loading magnitude (red=positive, blue=negative). (C) PC1-PC4 spatial representations along normalized body length (0=head, 1=tail). Each plot shows PC loading magnitude across body axis, capturing progressively finer spatial modes related to tail dynamics and local curvature patterns. (D) Mean PC eigencoefficients by developmental stage and adult condition. Bubble size: mean coefficient magnitude; color: sign (red=positive, blue=negative).

To examine the difference in eigenoikopleura coefficient features, we used a bubble grid chart, where the hue represents the sign (i.e. positive or negative coefficient) and radius represent the mean amplitude (Figure 3D). Thus, development reweights the same set of postural modes, with adults showing the broadest and most context-dependent PC usage. This analysis demonstrates that developmental stages and adult behavioral contexts occupy distinct regions of posture space, with adults exhibiting the greatest postural diversity.

To probe how these postural modes assemble into behaviors we employed a low-dimensional spatiotemporal embedding approach to reveal stereotyped behaviors exhibited by *Oikopleura* across post-embryonic development. For this approach, we used as input 49656 frames from a subset of balanced data across all three developmental stages. Unsupervised embedding resolved seven recurring clusters of behavior (Figure 4A) spanning quiescent to burst-like dynamics. All stages are represented in the single behavior space, but cluster occupancy is developmentally biased: larvae and juveniles emphasize lower-intensity regimes (clusters 1, 3 and 4) (Figure 4B and 4C) whereas adults disproportionately populate high-performance states (clusters 0 and 6) (Figure 4D). In adults, context (inside vs. outside the house) further reweights state usage. For example, clusters 0 and 4 are overrepresented in in house adults (relative to out of house adults), while clusters 1, 2 and 5 are overrepresented (Figure 4E and F). Only cluster 6 seems to be a novel state present in out of house adults but not in in house adults (or any other developmental stage) (Figure 4E and F). Overall, our analysis indicates repertoire conservation with developmental and contextual reweighting.

**Figure 4.**
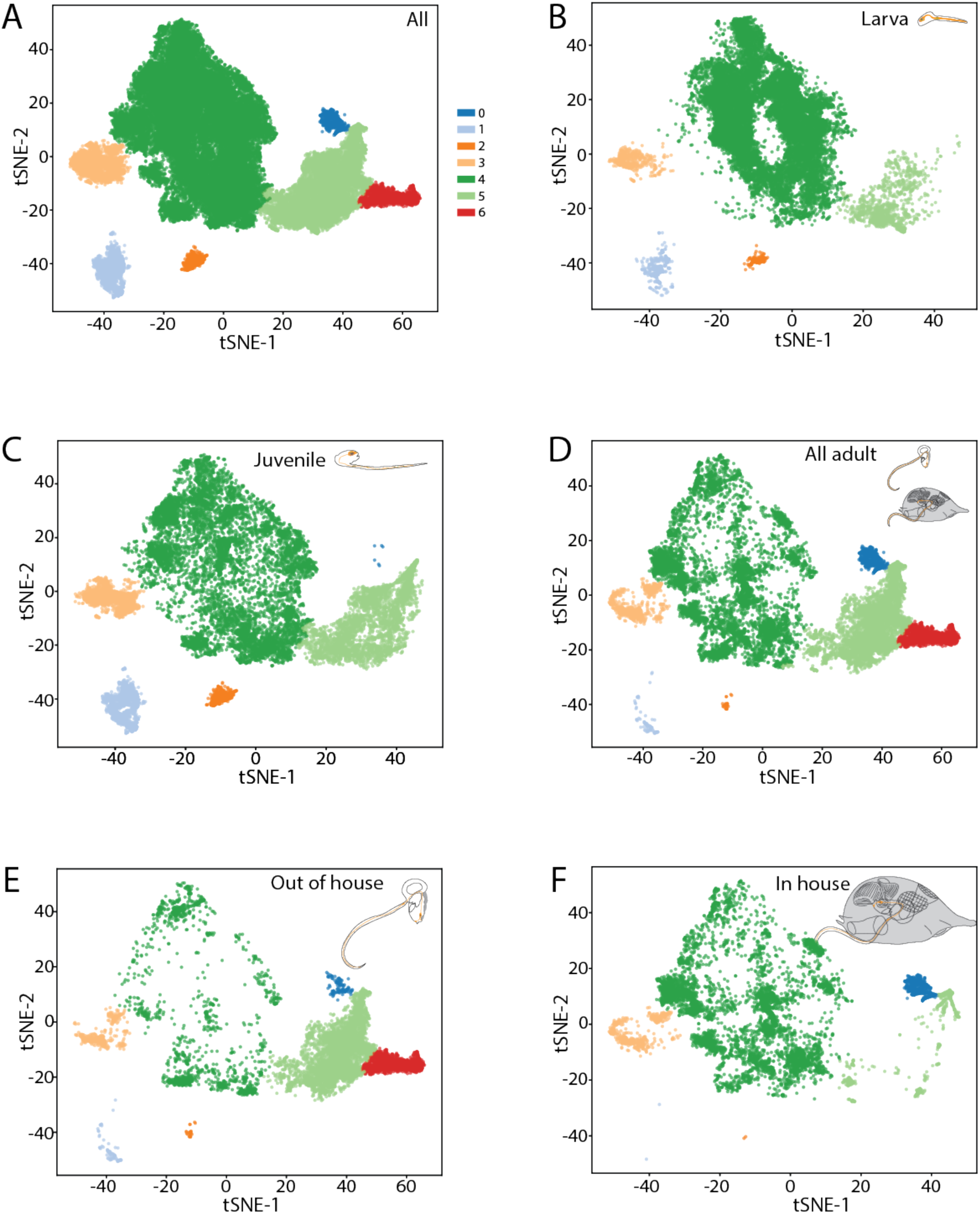
Unsupervised clustering reveals conserved behavioral modes across development. (A) t-SNE embedding of behavioral windows (n=45,872 windows from 57 individuals; 3 s windows, 1 s stride). Each point represents a behavioral window characterized by 8 features (velocity, acceleration, angular velocity, tail-beat frequency/amplitude, tortuosity, path complexity, MSD slope; see STAR Methods). Dimensionality reduction: features 7 PCs (95.3% variance), t-SNE (perplexity=750, stage-balanced training: n=15,290 per stage). Colors indicate HDBSCAN cluster assignment (7 states; parameters: min_cluster_size=25, min_samples=5, epsilon=0.15×median distance). Gray: unassigned noise points (2.3% of windows). (B–F) Stage-specific embeddings showing behavioral state distributions. (See also Figure S1)

To move from cluster identity to temporal structure, we fit Gaussian HMMs (full covariance; K by BIC) on the PC1-PC7 posture features. The emissions occupy distinct combinations of eigenoikopleuras (Figure 5A), and the resulting 13 hidden states provide a time-resolved scaffold over the seven unsupervised clusters. State usage is developmentally and contextually reweighted (Figure 5B-D): larvae concentrate in extended/low-activity states (S1), juveniles distribute occupancy toward moderate swimming (S4-S5 and S10, S11), and adults shift with context, inside the house favoring constrained, localized motion (e.g., S4/S10/S11) and outside the house favoring extended/propulsive states (e.g., S2/S9) and inside the house favoring constrained, localized motion (e.g., S4/S10/S11). States S4, S10, and S11 are shared across stages but with context-dependent shape realizations, suggesting behavioral homology despite morphological differences. Representative centerlines confirm distinct postural signatures (Figure 5E-G).

**Figure 5.**
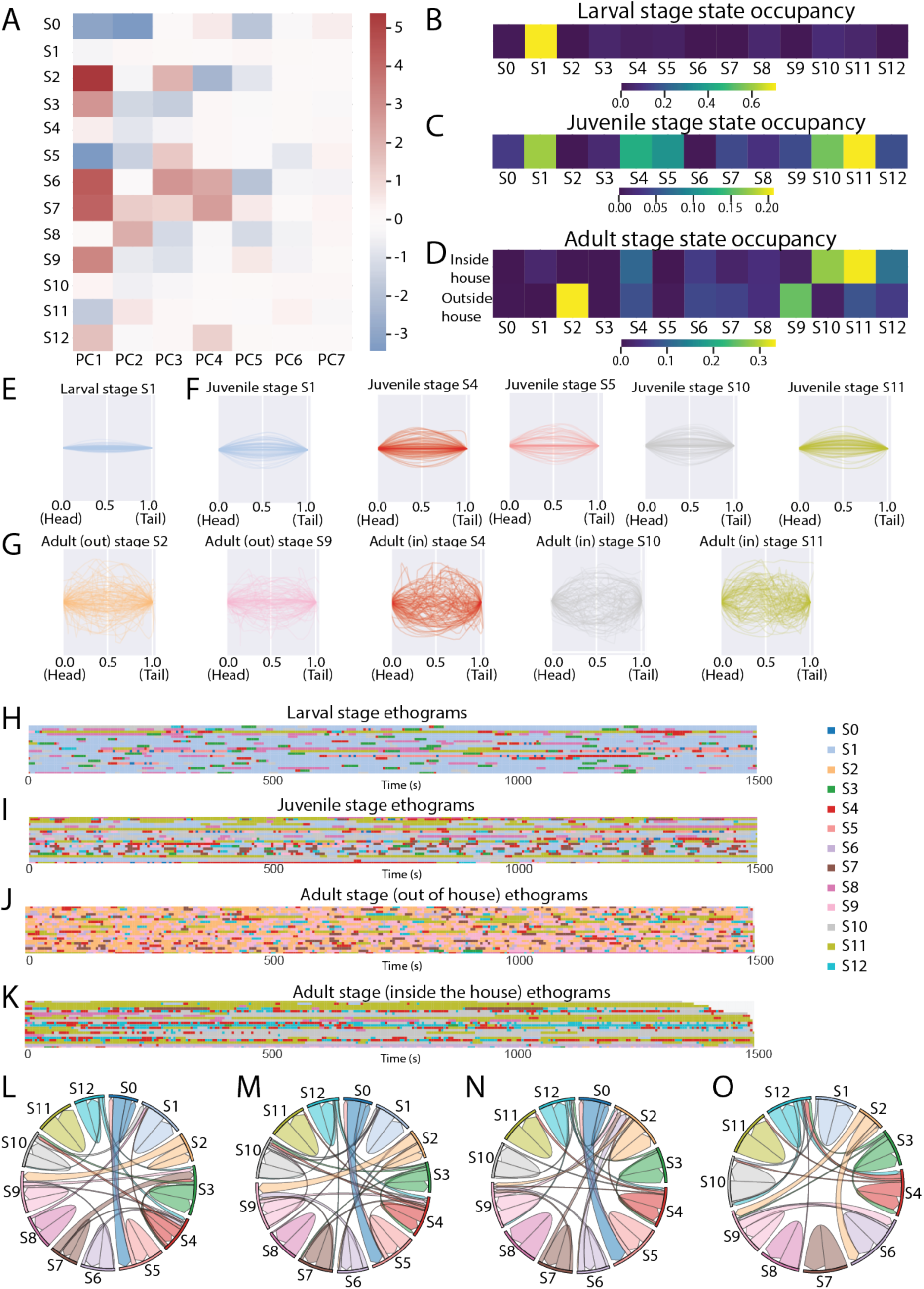
Hidden Markov model reveals temporal structure of behavioral sequences. (A) HMM emission means in PC space. Heatmap shows mean PC1-PC7 values for each of 13 hidden states (S0-S12) identified by Gaussian HMM (full covariance; K selected by BIC; see Methods). Rows: states; columns: PCs. Color: z-scored PC value (red=high, blue=low). States capture distinct combinations of postural modes. (B–D) State occupancy (fraction of frames) by developmental stage and adult condition. Color intensity: occupancy fraction. Each row sums to 1.0. (B) Larval stage. Dominated by states S0 (extended posture, 35% occupancy) and S1 (subtle undulation, 28%). Minimal occupancy of high-curvature states (S4, S11). (C) Juvenile stage. More distributed occupancy: S4 (moderate swimming, 21%), S5 (active tail-beating, 18%), with reduced S0 occupancy (15%) compared to larvae. (D) Adult stage occupancy by context. Inside house: S4 (32%), S10 (18%), S11 (15%) predominate states characterized by constrained, localized motion. Outside house: S2 (28%), S4 (22%), S9 (16%) states involving extended postures and propulsion. (E–G) Representative state-specific body shapes. Normalized centerlines (n=200 per state; gray=individuals, colored=mean) illustrate postural signatures. (E) Larval states: S1 (extended, minimal curvature); Juvenile S1 (slight anterior bend). (F) Juvenile states: S4 (active swimming, moderate tail bend), S5 (pronounced undulation), S10 (localized tail curvature), S11 (high-frequency tail motion with straight body). (G) Adult states: S2 (outside house, extended C-shape), S9 (outside, relaxed posture), S4 (inside, compact with tail wrap), S10 (inside, mid-body bend), S11 (inside, tight posterior coil). (H–K) Ethograms showing HMM state sequences over time (1,500 s sessions). Each row: one individual; colors: states S0-S12. (H) Larval stage. Prolonged state persistence (median dwell time S0=124 s, S1=89 s), reflecting limited behavioral repertoire. (I) Juvenile stage. Increased state diversity and transitions (median dwell times: S4=38 s, S5=29 s). (J) Adult outside house. Rapid state switching (median dwell S2=18 s) with episodic bursts of high-activity states.(K) Adult inside house. Complex temporal patterning with frequent transitions between S4, S10, S11 (median dwell times: 22 s, 31 s, 27 s respectively), reflecting house-maintenance behaviors. (L–O) State transition diagrams (chord plots). Arc thickness: transition probability; color: originating state. (L) Larval stage transitions. Strong self-transitions with limited interstate switching. (M) Juvenile stage transitions. Reduced self-persistence increased cross-state transitions. (N) Adult outside house. Highly connected network with S2 -S4, S4-S9, and S9-S2 as dominant transitions. (O) Adult inside house. Distinct transition structure: S4 - S10 and S10-S11form dominant cycles, reflecting stereotyped house-maintenance sequences. Model evaluation (held-out test set, 20% of sequences): training log-likelihood=-89,342; test log-likelihood=-22,105; BIC=179,284. Rare state pruning merged 1 state (<1% occupancy) into nearest neighbor (see Methods). Final model: K=13 states, median state dwell time=27 s (range: 8-124 s across stages/contexts). (See also Figure S2)

The HMM also exposes sequencing (Figure 5H-K): larvae show long dwell times and strong self-persistence; juveniles break into more frequent transitions; adults exhibit rapid switching outside the house and stereotyped maintenance cycles inside (S4-S10-S11). Transition graphs formalize these patterns (Figure 5L-O), with S4/S10/S11 acting as hubs. Together, these results show that development and adult context reshape how conserved behaviors are chained in time-altering dwell times and transition backbones-rather than introducing entirely new states.

## DISCUSSION

In this study we have used high throughput tracking and multi-dimensional behavioral representations to characterize changes in the behavioral repertoire of *Oikopleura* throughout ontogeny. Our work bridges the gap between invertebrates and vertebrates in terms of behavioral developmental plasticity.

Comparative studies in non-limbed chordates suggest a developmental sequence of movement culminating in adult swimming patterns^36^. Our observations in *Oikopleura* support this hypothesis: larvae and juveniles exhibit whole-body bends, which transition to undulatory swimming in adults. Behavioral developmental plasticity has been well documented in aquatic vertebrate models such as zebrafish^37,38^ and Amazon mollies^8^. In zebrafish, studies report increased locomotor activity with developmental progression^37,38^, and one study also observed heightened thigmotaxis^37^, akin to what we found in *Oikopleura*. Amazon mollies exhibit increased behavioral plasticity for several weeks, followed by a decline^8^. In zebrafish, plasticity is not limited to spontaneous locomotion; complex behaviors like predation and collective behavior also intensify during development^39,40^. Whether *Oikopleura* behaviors beyond swimming and house-related activities show similar plasticity remains to be explored.

Insects provide additional context with respect to behavioral developmental plasticity amongst invertebrates. In *Drosophila melanogaster*, overall behavioral complexity and stereotypy remain stable across adulthood, despite repertoire changes^41^. Honeybees also exhibit age-related behavioral shifts driven by physiological development, environmental factors, and social interactions^42–45^, a phenomenon recently analyzed using multi-dimensional approaches that revealed differences in movement characteristics and transition dynamics^46^. These findings mirror our observations in *Oikopleura dioica*, suggesting that behavioral developmental plasticity may affect common behavioral features across species even if they are evolutionarily distant.

We have shown that *Oikopleura* modulates swimming kinematics, area exploration, and motor module usage and transitions between modules depending on developmental stage and context (inside vs. outside the house). The molecular and neural mechanisms underlying this plasticity remain poorly understood. Across species, changes in swimming patterns have been linked to diverse mechanisms such as context-dependent sensory inputs, dynamic state shifts, and neuromodulatory control^47–49^. In *Oikopleura*, dopamine has been implicated in regulating swimming and idling outside the house^50^, but its role in house inflation and filter feeding is unknown. Although we did not identify molecular or cellular drivers of plasticity, prior work on *Oikopleura*’s nervous system provides clues. Mature neuromuscular synapses appear around 15 hours (juvenile stage) and peak in day-3 adults^10^ potentially correlating with the transition from simple tail bends to rhythmic sinusoidal swimming. While neuron numbers remain stable across development, support cells in the caudal nerve cord increase dramatically, suggesting a role in motor modulation^10^. Their molecular identity remains unknown, but proximity to motor neurons implies functional significance. Future studies can leverage established tools in *Oikopleura* and related tunicates (e.g., CRISPR/Cas9 genome editing^51^, calcium imaging, chemogenetics^52–55^ ) to uncover the molecular and neuronal basis of behavioral plasticity.

While we have characterized behavioral changes across *Oikopleura* ontogeny, an important open question is how early-life environmental stress (reviewed by Ehlman et al. ^56^) impacts behavioral developmental plasticity during later stages of ontogeny. Most research on stress effects has focused on terrestrial organisms, leaving marine species largely unexplored. *Oikopleura dioica*, one of the most abundant planktonic species^57,58^, is highly sensitive to anthropogenic stressors^59,60^. Combining ontogenetic tracking with early-life stress exposure could reveal mechanisms linking early experiences to behavioral outcomes later in life.

More generally, despite its miniaturized nervous system (∼130 neurons^9–11^, the smallest documented in chordates), *Oikopleura* exhibits surprising behavioral complexity. This suggests an evolutionary limit beyond which further neuronal reduction would compromise function. Its circuits may represent minimal units below which complex behaviors would like not be possible.

## RESOURCE AVAILABILITY

### Lead contact

Further information and requests for resources and reagents should be directed to and will be fulfilled by the lead contact, M. Chatzigeorgiou

### Materials availability

This study did not generate any new materials.

### Data and code availability

- Raw video data have been deposited in Zenodo. The DOIs are:

https://doi.org/10.5281/zenodo.17502188
https://doi.org/10.5281/zenodo.17464322
https://doi.org/10.5281/zenodo.17455742
https://doi.org/10.5281/zenodo.17503716
- The DeepLabCut based DNN trained to recognize landmarks on Oikopleura dioica larvae, juveniles and adults has been deposited in Zenodo together with the training dataset. The DOIs are:

https://doi.org/10.5281/zenodo.17473173
https://doi.org/10.5281/zenodo.17459389
https://doi.org/10.5281/zenodo.17491493
- All original code will be deposited at Zenodo and will be publicly available as of the date of publication. Code to analyze behavioral data, is already available here:

https://github.com/ChatzigeorgiouGroup/Atlas_behavior_Oikopleura
- Any additional information required to reanalyze the data reported in this paper will be made available from the lead contact upon request.

## ACKNOWLEDGMENTS

We would like to thank members of the Chatzigeorgiou lab for valuable feedback on the manuscript. We acknowledge funding from the Michael Sars Centre core budget (NFR grant 234817 “Sars International Centre for Marine Molecular Biology Research, 2013–2022”) and grants from the Research Council of Norway to M.C. (339399 and 335582) and D.C. (250005).

## AUTHOR CONTRIBUTIONS

Conceptualization: O.T., S.N., R.S.M.B., M.C.; Methodology: O.T., S.N., R.S.M.B., M.C.; Data curation: O.T., R.S.M.B., S.N.; Investigation: O.T., R.S.M.B., S.N.; Visualization: O.T., R.S.M.B., S.N.; Resources: M.C., D.C., A.E.A; Funding acquisition: M.C., D.C.; Project administration: M.C.; Supervision: M.C.; Writing – original draft: O.T., S.N., R.S.M.B., M.C.; Writing – review and editing: O.T., S.N. R.S.M.B., ,D.C., A.E.A, M.C.

## SUPPLEMENTAL INFORMATION

**Figure S1: HMM transition networks and state characteristics**

(A-D) State transition probability matrices for larvae (A), juveniles (B), adults outside (C), and inside house (D). Diagonal = self-persistence; off-diagonal = interstate transitions. (E-H) State emission means in PC space for each condition. (I-L) Representative state-specific centerline shapes showing postural signatures. Developmental progression shows: decreased state persistence (dwell times: 124s to 27s), increased transition complexity, and context-dependent transition networks in adults.

**Figure S2 Behavioral cluster assignments over time for individuals across developmental stages and adult contexts.** (A-B) Larvae. (C-D) Juveniles. (E-F) Adults outside house. (G) Adults inside house. (H) Inside-house. White gaps in ethograms indicate tracking interruptions. Panels B, D F, H show normalized centerline overlays. Black line= mean, grey line= individual animal.

**Movie S1** Video showing Oikopleura dioica larvae swimming in the behavioral arena.

**Movie S2** Video showing Oikopleura dioica juveniles swimming in the behavioral arena.

**Movie S3** Video showing an Oikopleura dioica adult freely-swimming in the behavioral arena.

**Movie S4** Video showing an Oikopleura dioica adult inside the house filter feeding. The animal is located inside the behavioral arena.

## STAR★METHODS

### KEY RESOURCES TABLE

| REAGENT or RESOURCE | SOURCE | IDENTIFIER |
| --- | --- | --- |
| **Antibodies** |  |  |
| **Bacterial and virus strains** |  |  |
| **Biological samples** |  |  |
| **Chemicals, peptides, and recombinant proteins** |  |  |
| **Critical commercial assays** |  |  |
| **Deposited data** |  |  |
| Raw videos of *Oikopleura dioica* (larvae, juveniles, adults inside or outside the house) | https://doi.org/10.5281/zenodo.17502188<br>https://doi.org/10.5281/zenodo.17464322<br>https://doi.org/10.5281/zenodo.17455742<br>https://doi.org/10.5281/zenodo.17503716 |  |
| DLC DNN trained to recognize *Oikopleura dioica* body parts and training data | https://doi.org/10.5281/zenodo.17473173<br>https://doi.org/10.5281/zenodo.17459389<br>https://doi.org/10.5281/zenodo.17491493 |  |
| **Experimental models: Cell lines** |  |  |
| **Experimental models: Organisms/strains** |  |  |
| *Oikopleura dioica* | Bergen, Vestland, Norway | N/A |
| **Oligonucleotides** |  |  |
| **Recombinant DNA** |  |  |
| **Software and algorithms** |  |  |
| DeepLabCut | Lauer et al. <sup>33</sup> |  |
| Matplotlib | https://github.com/matplotlib/matplotlib | Version 3.10.8 |
| Numpy | https://github.com/numpy/numpy | Version 2.3.5 |
| scikit-learn | https://github.com/scikit-learn/scikit-learn | Version 1.5 |
| SciPy | https://github.com/scipy/scipy | Version 1.16.3 |
| pandas | https://github.com/pandas-dev/pandas | Version 2.3.3 |
| Seaborn | https://github.com/mwaskom/seaborn | Version 0.13.2 |
| openTSNE | https://github.com/pavlin-policar/openTSNE | Version 1.0.4 |
| hmmlearn | https://github.com/hmmlearn/hmmlearn | Version 0.3.3 |
| HDBSCAN | https://github.com/scikit-learn-contrib/hdbscan | Version 0.8.39 |
| **Other** |  |  |

### EXPERIMENTAL MODEL AND STUDY PARTICIPANT DETAILS

Wild *Oikopleura* were collected in coastal waters near Bergen, Norway, and cultivated for several generations at the Michael Sars Centre, University of Bergen, Norway, before use. The animals were kept according to standard protocol at 12±0.5 °C resulting in a 7-day life cycle^61^.

### METHOD DETAILS

#### *Oikopleura dioica* behavioural recording

For the tracking of the *Oikopleura* in its different stages, a customized behaviour set up containing Nikon SMZ1500 stereomicroscope fitted with a HR Plan Apo 1x (N/A 0.131/WD54 mm) as previously described^13,32^. The animals were recorded in 0.8% agarose arenas (10 mm diameter and 3 mm high, approximately 236 mm^3^) containing filtered sea water (FSW), at 30 frames/second speed.

For the larval and juvenile stages, controlled fertilizations were performed. Eggs from naturally spawned females were fertilized with sperm pool obtained from a minimum of 3 to 4 males. The first cell division around 20-30 minutes after fertilization was checked to confirm the fertilization and the embryos were kept at 13 °C until recording. For larval stage, fertilized eggs were incubated until 7-8 hours post fertilization (hpf) and larvae were recorded for a total of 10 minutes. For the juvenile stage, animals were incubated until 16-18 hpf and recorded for a total of 10 minutes.

For the adult stage, animals were recorded on day 5 for a total of 10 minutes. The first 5 minutes were recorded inside their houses (marked by the addition of *Synechococcus* cyanobacteria) and forced out of the house and recorded for additional 5 minutesto register both of their behavioural states (inside and outside their houses).

#### DeepLabCut model development for tracking of *Oikopleura dioica* in different developmental stages

To evaluate the behaviour of the animals on the recordings obtained, three different models (larvae, juvenile and adult models) were developed using machine learning software Deeplabcut (DLC) 3.0.0rc6, training the AI tool to detect and track the larvae, juvenile and adult animals. For the training of the models, 15-16 videos were manually annotated, with a minimum of 20 randomly selected frames per video.

#### Data processing

Behavioral tracking data were obtained using DeepLabCut (v3.0.0rc6) trained on manually annotated frames spanning all developmental stages (larva, juvenile, adult). Tracking confidence was filtered at likelihood ≥0.99, and statistical outliers were removed using the interquartile range method (IQR threshold = 2.0). Teleportation artifacts (frame-to-frame displacement >100 pixels) were eliminated. To ensure temporal continuity, only tracking segments ≥3 s (90 frames at 30 fps) was retained for larval and juvenile stages; adult stage segments required ≥1 s continuity. Internal gaps in tracking were filled using linear interpolation when bracketed by valid detections. Coordinates were temporally downsampled 1:5 (6 Hz) for computational efficiency while preserving behavioral dynamics.

#### Behavioral metrics extraction

Postural metrics: Body centrelines were extracted by interpolating through trunk and tail anchor points using piecewise cubic Hermite interpolation (PCHIP) to 21 evenly spaced points. Centrelines were normalized by translating the head to the origin, rotating the tail onto the positive x-axis, and scaling total length to unity. Local curvature was computed using Savitzky-Golay filtering (window=15 frames, polynomial order=2) of centreline derivatives. Body axis orientation (theta) was defined as the angle from mean head position to mean tail position.

Kinematic metrics: Center of mass (COM) was calculated as the mean position of trunk markers. Instantaneous velocity was computed as the Euclidean displacement of COM between consecutive frames, converted to µm/s using a calibrated pixel-to-micrometre ratio (11.56 µm/pixel). Angular velocity (omega) was derived by unwrapping body axis angles within continuous tracking segments and applying finite differences, preventing NaN propagation across gaps. Acceleration was computed as the temporal derivative of velocity.

Tail-beat analysis: Tail-beat frequency was determined at native temporal resolution (30 fps) using fast Fourier transform (FFT) of the lateral tail displacement signal with Hann windowing. Dominant frequency was refined via quadratic interpolation of the peak spectral bin. Tail-beat amplitude was quantified as half the peak-to-peak displacement over 1 s windows.

Window-based features: Behavioral metrics were aggregated over sliding 3 s windows with 1 s stride. For each window, we computed: (1) 95th percentile velocity and acceleration, (2) fraction of frames above a movement threshold (defined as 3× median absolute deviation of per-individual velocity), (3) path tortuosity (path length / net displacement), (4) path complexity (entropy of normalized x-y correlation), and (5) mean-squared displacement (MSD) slope over short lags (1-3 frames).

#### Unsupervised behavioral state discovery

Feature preprocessing: Eight features were selected for behavioral state classification: tortuosity, path complexity, MSD slope, tail-beat frequency, tail-beat amplitude, velocity (95th percentile), acceleration (95th percentile), and angular velocity (95th percentile). Positive-valued features (velocity, acceleration, angular velocity, tail-beat amplitude) were log₁₊-transformed. Features were robustly clipped to the 0.5-99.5th percentile range and median-centered within individuals to account for inter-individual variability. Global standardization (z-score) was then applied across the dataset.

#### Dimensionality reduction and embedding

Preprocessed features were reduced to 7 principal components (PCs) using PCA, capturing >95% cumulative explained variance. To ensure balanced representation across developmental stages, data were downsampled such that each stage (larva, juvenile, adult) contributed equally to the training set (n = [minimum stage count] per stage). The balanced dataset was embedded in 2D using t-distributed stochastic neighbor embedding (t-SNE) via openTSNE with perplexity=750 (affinity: approximate nearest neighbors; initialization: PCA; early exaggeration iterations=250, total iterations=1000).

Clustering: Behavioral states were identified using Hierarchical Density-Based Spatial Clustering of Applications with Noise (HDBSCAN). To determine optimal hyperparameters, we performed a grid search over min_cluster_size (15-30), min_samples (None, 1, 5), and epsilon (0-20% of median pairwise distance in embedding space), evaluating cluster stability and noise fraction. The resulting clustering yielded K=7 distinct behavioral states. Cluster labels from the balanced embedding were propagated to the full dataset using k-nearest neighbours (k=7, distance weighting) in the t-SNE embedding space.

#### Hidden Markov model analysis

Sequence preparation: Individual animal trajectories in PC space (7 dimensions) were treated as sequences, with minimum sequence length 30 frames. Sequences were split 80:20 into training and held-out test sets.

Model fitting. Gaussian Hidden Markov Models (HMMs) were fit with K=[5-15] hidden states using the hmmlearn package. For each K, models were initialized via K-means clustering of the training data to set emission means and covariances (full covariance matrices). Start probabilities were derived from cluster occupancy, and transition matrices were initialized with strong self-transition bias (0.8 diagonal). Expectation-maximization (EM) was run for up to 500 iterations (tolerance=10⁻⁴) from 8 random initializations, retaining the model with highest training log-likelihood.

Model selection: Models were compared using Bayesian Information Criterion (BIC = -2·LL + p·log(N), where p = number of free parameters, N = total training frames) and held-out log-likelihood. The selected model (K=14) balanced parsimony and explanatory power.

State pruning: States occupying <1% of total frames were identified as rare and merged into the nearest state by Euclidean distance in emission mean space. The model was then refit with the reduced state count (K_final=13).

Characterization: Viterbi decoding was applied to all sequences to infer the most likely state sequence. State dwell times (run lengths) were computed. State occupancy and transition probabilities were computed separately for each stage × experimental condition combination.

#### Shape Analysis

For each behavioral state, normalized centerlines (n=[min 100, max 500] per state) were sampled from time windows assigned to that state. Quality filtering removed centerlines with lateral deviation >0.6 body lengths or adjacent segment angles >2.0 radians (∼115°). Mean centerline shapes were computed per state and visualized to reveal state-specific postural signatures.

#### Statistical Comparisons

Between-stage comparisons. Behavioral metrics were averaged per individual (across all time windows), yielding one value per animal. For comparisons across three developmental stages, Kruskal-Wallis H-tests were performed. Post-hoc pairwise comparisons used Mann-Whitney U tests with Holm-Bonferroni correction for multiple testing. For two-group comparisons, Mann-Whitney U tests were applied directly.

#### Sample sizes

Analyses included 159 larvae (26.5 h of tracking), 106 juveniles (17.7 h), and 60 adults (10.0 h). After quality control, filtering, resampling the dataset comprised 572,559 frames for larvae, 381,706 frames for juveniles, and 216,060 frames for adults (1,170,325 frames in total, 54.2 h of behavior). A balanced subset of 49,656 frames (2.5 h) was used for clustering analyses. Significance and reporting. Statistical significance was assessed at α=0.05 after correction for multiple comparisons where applicable. All tests were two-tailed. Data are presented as median ± interquartile range unless otherwise noted. Exact p-values are reported in figure legends; p<0.001 is indicated as such.

