## Supplementary figures and images for "Behavioral ontogeny in a pelagic tunicate reveals the deep origins of chordate behavioral developmental plasticity"

### Figure S1

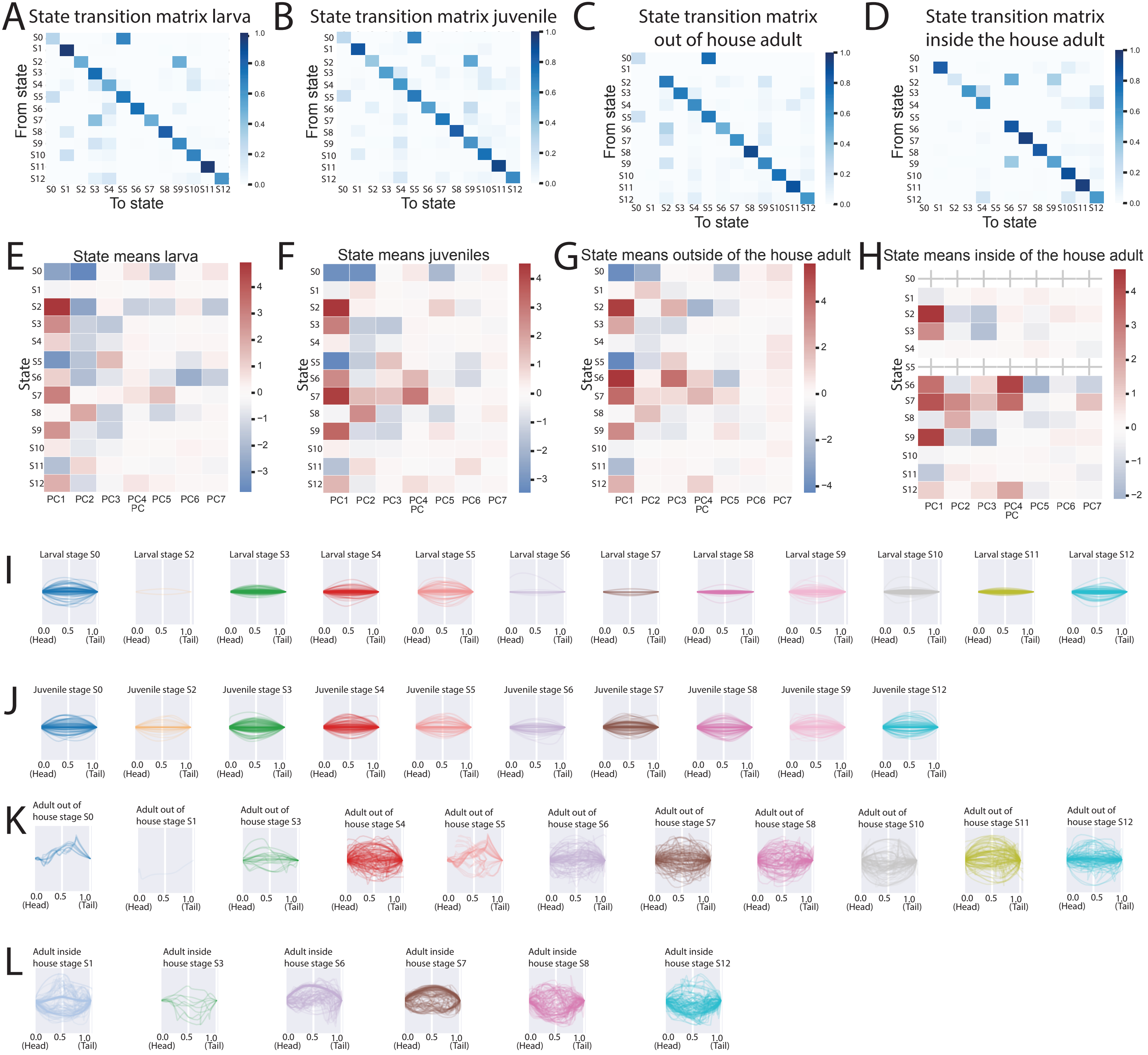

### Figure S2

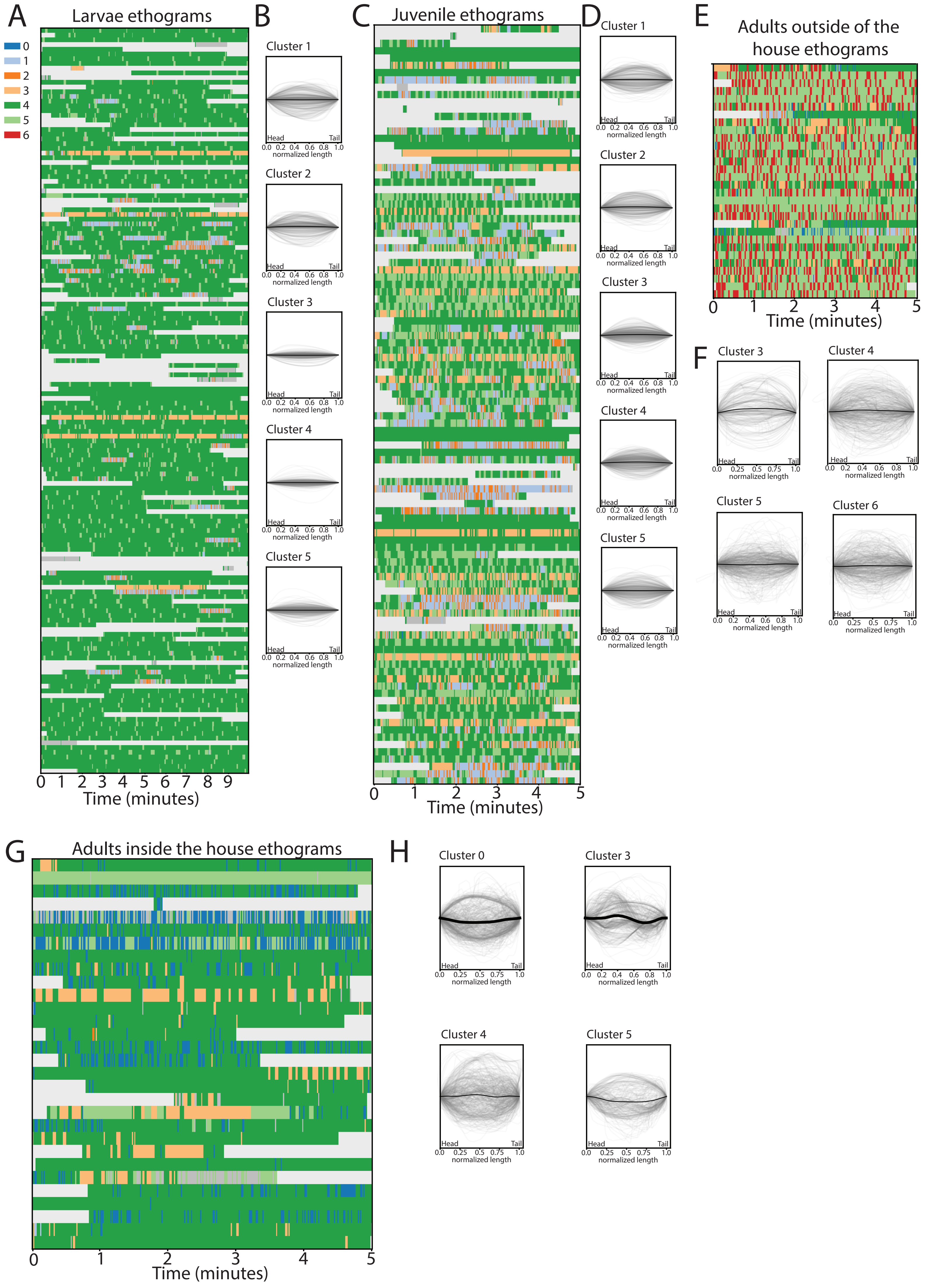
